# Integrated multi-cohort analysis of the Parkinson’s disease gut metagenome

**DOI:** 10.1101/2022.07.20.500694

**Authors:** Joseph C. Boktor, Gil Sharon, Leo A. Verhagen Metman, Deborah A. Hall, Phillip A. Engen, Zoe Zreloff, Daniel J. Hakim, John W. Bostick, James Ousey, Danielle Lange, Gregory Humphrey, Gail Ackermann, Martha Carlin, Rob Knight, Ali Keshavarzian, Sarkis K. Mazmanian

## Abstract

**Background:** The gut microbiome is altered in several neurologic disorders including Parkinson’s disease (PD).

**Objectives:** Profile the fecal gut metagenome in PD for alterations in microbial composition, taxon abundance, metabolic pathways, and microbial gene products, and their relationship with disease progression.

**Methods:** Shotgun metagenomic sequencing was conducted on 244 stool donors from two independent cohorts in the United States, including individuals with PD (n=48, n=47, respectively), environmental Household Controls (HC, n=29, n=30), and community Population Controls (PC, n=41, n=49). Microbial features consistently altered in PD compared to HC and PC subjects were identified. Data were cross-referenced to public metagenomic datasets from two previous studies in Germany and China to determine generalizable microbiome features.

**Results:** The gut microbiome in PD shows significant alterations in community composition. Robust taxonomic alterations include depletion of putative “beneficial” gut commensals *Faecalibacterium prausnitzii* and *Eubacterium* and *Roseburia* species, and increased abundance of *Akkermansia muciniphila* and *Bifidobacterium* species. Pathway enrichment analysis and metabolic potential, constructed from microbial gene abundance, revealed disruptions in microbial carbohydrate and lipid metabolism and increased amino acid and nucleotide metabolism. These global gene-level signatures indicate an increased response to oxidative stress, decreased cellular growth and microbial motility, and disrupted inter-community signaling.

**Conclusions:** A metagenomic meta-analysis of PD shows consistent and novel alterations in taxonomic representation, functional metabolic potential, and microbial gene abundance across four independent studies from three continents. These data reveal stereotypic changes in the gut microbiome are a consistent feature of PD, highlighting potential diagnostic and therapeutic avenues for future research.

## INTRODUCTION

Parkinson’s Disease (PD) is a progressive movement disorder estimated to affect over 1.2 million people in the United States by 2030^1–3^. Only 15% of diagnoses are attributed to a monogenic cause, suggesting a strong environmental role in PD^4^. Indeed, environmental risk factors for PD include geographic location, exposure to pesticides, and certain solvents and metals^5–7^. PD presents with motor symptoms including tremors, bradykinesia, muscle rigidity, impaired posture, and difficulty in speech and swallowing^8^. Curiously, non-motor symptoms, including hyposmia, depression, sleep disruption, and gastrointestinal (GI) distress such as constipation may appear years prior to PD diagnosis^9–12^.

The gut microbiome represents hundreds of species of bacteria, fungi, archaea, and viruses that have been linked to the regulation of the immune, metabolic, and nervous systems^13–17^.

Dysbiosis, defined as a comparative alteration in microbiome composition^18^, has been implicated in PD and may be linked to the comorbidity of prodromal constipation^19–30^. Previous studies analyzing the fecal microbiome in PD via 16S amplicon profiling have found changes in multiple taxa compared to healthy controls. In contrast to 16S rRNA amplicon sequencing, shotgun metagenomics provides an untargeted sequencing approach that allows for the estimation of both microbial community composition and its potential functions^31,32^. Notable depletions at the genus level in PD are the commensal bacteria *Roseburia, Lachnospiraceae, Blautia, Prevotella, Faecalibacterium*, and *Eubacterium*, while *Lactobacillus, Bifidobacterium, Akkermansia*, and *Alistipes* are typically enriched in PD compared to healthy controls^23,25,27,33–36^. Studies examining the gut metagenome in PD have identified PD-related alterations in microbial metabolic pathways including homocysteine, folate, and sulfur metabolism^20,36–38^. Using integrated modeling of metagenomic and metabolomic data, these studies have identified differentially abundant plasma metabolites in PD that are consistent with dysregulation in microbial metabolic pathways including cysteine metabolism, a process associated with oxidative stress^37^.

While prior work has focused on PD-associated microbial dysbiosis at the taxonomic level, it is still unknown how these taxonomic differences translate to functional contributions in the development of PD. Accordingly, we sought to characterize alterations of gut microbiome metabolic potential in PD in both the early and late stages of disease, incorporating household control samples that allow insights into the shared environment between people discordant for disease. We conducted metagenomic sequencing on stool samples of 244 participants across two distinct cohorts in the United States and extended our findings with an additional 137 samples from publicly available datasets from two additional cohorts in Germany and China. This study represents the first meta-analysis of metagenomic data in PD, integrating findings across populations in three continents to identify novel microbial features consistently altered in PD, revealing that globally generalizable signatures in the gut are associated with a major neurogenerative disorder.

## METHODS

### Study Design

Samples were provided by the BioCollective (TBC) and Rush University Medical Center (RUMC; Chicago, IL) with respective recruitment of: individuals with Parkinson’s Disease (PD, n=48, n=47), Household Controls (HC, n=29, n=30), and healthy Population Controls (PC, n=41, n=49). RUMC participants were recruited at the Movement Disorders Clinic with clinical assessments conducted at the RUMC Parkinson’s Disease Gastroenterology Clinic (PDGC).

### RUMC Participants

Prior to the clinical visit, movement disorder specialists examined and confirmed the diagnosis of all PD participants at a baseline screening. Parkinsonian symptoms were assessed using the Unified Parkinson’s Disease Rating Scale (UPDRS)^39^ and Hoehn and Yahr (H&Y) staging scale^40^.

Inclusion criteria for PD participants: 1) age between 40 and 80 years, and 2) a current diagnosis of PD (UK Brain Bank Criteria, H&Y stages 1-4 inclusive)^41^. Inclusion criteria for HC and PC participants: 1) age between 40 and 80 years, 2) no history of neurological disorders or neurodegenerative disease, and 3) for HC only -living in the same household and consuming a similar diet as an enrolled PD participant. Exclusion criteria for PD, HC and PC participants: 1) Presence of symptomatically active gastrointestinal diseases such as inflammatory bowel disease (IBD) or celiac disease (except for hemorrhoids, hiatal hernia, or occasional (˂3 times a week) heartburn), 2) Antibiotics, probiotics (except yogurt), prebiotics usage, or intentional diet change within 12 weeks of sample collection, 3) Abdominal surgeries for GI disease such as bowel resection, diverticular surgery, colostomy (surgery for hemorrhoids and cholecystectomy or appendectomy for benign disease more than 5 years prior to enrollment were allowed), 4) Symptomatic functional GI disease that could impair intestinal motility such as scleroderma, 5) Acute illness requiring hospitalization, 6) Pre-existent organ failure or comorbidities: a) liver disease (cirrhosis or persistently abnormal AST or ALT that are 2X˃ normal); b) kidney disease (creatinine ˃ 2.0mg/dL); c) uncontrolled psychiatric illness; d) clinically active lung disease or decompensated heart failure; e) known HIV infection; f) alcoholism; g) transplant recipients; h) diabetes, 6) Presence of short bowel syndrome or severe malnutrition, 7) Chronic use of diuretics, 8) Chronic use of NSAIDS. A washout period of three weeks was needed before the subject could be enrolled into the study. Low dose aspirin was allowed. Participants from Rush University signed the RUMC Institutional Review Board (IRB) approved informed consent forms (ORA # 16111903 and 10062805) and the study was registered (ClinicalTrials.gov Identifier: NCT03705520).

### TBC Participants

Samples included from TBC’s biobank are approved under guidance of the Caltech IRB. PD participants were self-identified, verifying that they have received a PD diagnosis from a qualified physician. HC criteria required only that they share a household with the PD participant, while PC patients were selected to match the PD sample averages for age, sex, BMI, and race and have no current illness.

### Sample Collection

Stool samples were self-collected at home using either the anaerobic home collection kit (BD Gaspak, Becton Dickinson and Company, Sparks, MD) or the BioCollector™ kit to minimize the exposure of stool to oxygen^42,43^. Upon arrival, Bristol stool scores were recorded, and samples aliquoted and stored at -80°C.

### Questionnaire Collection

Prior to fecal collection, RUMC PD and HC participants completed questionnaires regarding diet, smell, sleep and adverse events related to gastrointestinal symptoms. Dietary questionnaires included the automated self-administered 24-hour recall (ASA24®)^44^, and three month-recall Vioscreen Food and Frequency of Consumption Questionnaire (FFQ)^45^. Smell assessments used the University of Pennsylvania Smell Identification Test (UPSIT)^46^. Sleep evaluations used the Idiopathic Rapid Eye Movement (REM) Sleep Behavior Disorder (RBD) single-question screen (RBD1Q)^47^, the Pittsburgh Sleep Quality Index (PSQI) self-reported questionnaire to assess the sleep habits^48^, and the Munich ChronoType Questionnaire (MCTQ) pertaining to sleep, activity times, and jet lag^49^. Gastrointestinal evaluation used the Patient-Reported Outcomes Measurement Information System (PROMIS) gastrointestinal symptom scale to examine GI symptoms and severity^50^. TBC participants completed an optional TBC specific questionnaire documenting dietary habits, supplement usage, general and PD medication usage, demographic, and various miscellaneous metadata (Fig S1B). Additionally, PD participants from TBC were invited to complete the self-reported sections of the MDS-UPDRS (parts I & II). Further details about questionnaires are described in the supplementary material.

### Sequencing, Data Handling, and Pre-Processing

gDNA was extracted using the Qiagen MagAttract PowerSoil gDNA kit. DNA quality was evaluated visually via gel electrophoresis and quantified using a Qubit 3.0 fluorometer (Thermo-Fisher, Waltham, MA, USA) and Quant-iT PicoGreen dsDNA Assay Kit (Invitrogen, Waltham, MA, USA) for TBC and RUMC samples, respectively. Libraries were prepared using an Illumina Nextera XT library preparation kit following the standard protocol (Illumina, San Diego, CA, USA).

Samples were sequenced using 150 bp paired-end reads with the Illumina NextSeq and NovaSeq for TBC and RUMC samples, respectively. Taxonomic and functional profiles were generated with the bioBakery meta’omics workflow (v3.0.0-α.4)^31^. Metagenomic reads were filtered using KneadData (v0.7.4) to remove reads with low quality or that map to the human genome. Taxonomic profiles were generated with MetaPhlAn3 (v3.0.7) and functional profiling with HUMAnN3 (v3.0.0.α.3) mapping to the UniRef90 catalogue (UniRef release 2019_01).

UniRef90 relative abundance tables were then regrouped into the following higher-level organizations: Enzymes, MetaCyc pathways, Gene Ontology (GO), KEGG Orthology, protein families (Pfams), and eggNOGs. For each organization level, unmapped or ungrouped abundance variables were removed prior to analysis. A threshold of 1 million quality reads was set for study inclusion. In total, 227 samples, including 51 household and PD pairs, 83 healthy controls, and 88 PD samples met all listed criteria and were included for analysis.

### Community Composition Analysis

For community-level analyses, data were first rarefied to an even depth by generating pseudo-counts by multiplying MetaPhlAn3/HUMANn3 relative abundance profiles with total quality reads per sample and sub-sampling counts to match the minimum number of reads of a sample (> 1 million). Alpha-diversity metrics (i.e., observed features, Shannon’s Diversity Index, and Simpson’s Evenness) were calculated from MetaPhlAn3 taxonomic profiles using the microbiome R package v1.14.10. Statistical analysis was conducted using the mixed models specified below with lmer (lme4 R package v1.1-27.1) (Eq. 1; Eq. 2). Beta-diversity analysis was conducted using Aitchison distance calculated with the phyloseq R package (v1.36.0). To estimate the contribution of metadata variables on community composition, a Permutational Multivariate Analysis of Variance (PERMANOVA) was conducted using adonis (vegan package R v2.5-7) on the Aitchison distance of species abundance profiles with 99,999 permutations.

Each metadata variable was uniquely processed to remove samples with missing values and run in an independent model. Variables for usage of PD medications were tested exclusively within PD samples, removing the potential for donor group signatures confounding PD-specific variables. We then applied a *P*-value correction False Discovery Rate (FDR) for multiple comparisons using the Benjamini-Hochberg method.

### Statistical Analysis

All statistical analysis was conducted in R (v4.1.0). To determine feature associations with PD status, we utilized Multivariable Association with Linear Models (MaAsLin2)^51^. Prior to differential abundance testing, data tables were filtered for features with a minimum prevalence of 10%. Biobakery relative abundance tables were scaled to a total sum of one followed by an arcsine square root transformation for variance stability. We then applied a feature-level specific variance filter based on the variance distribution and the number of features present at each level. Two separate analyses were conducted in parallel for each feature level. The first includes all PD donors and all PC donors:

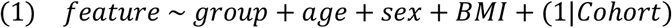

The second model includes all HC pairs and their respective PD donors:

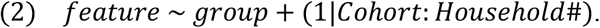

For our generalized mixed linear models, we denote (q ≤ 0.1) as significance and (q ≤ 0.25) as association.

As an assessment for the confounding potential of covariates, we devised an iterative testing strategy where relevant variables are selected and appended to a generalized linear model containing all participants with the following formula:

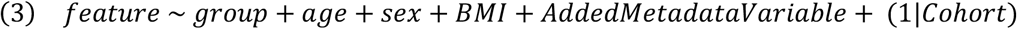

repeated for each feature level. The following variables on supplement intake or usage are tested for their overlap with disease signature feature significance: calcium, vitamin C, non-steroidal anti-inflammatories, proton pump inhibitors, laxatives. The impact of PD medications was similarly tested using a model akin to Eq.3, but only within PD donors and removing the donor group variable from the model. This strategy is designed to maximize use of provided metadata, whereas an analysis combining multiple metadata variables with sporadic responses in a single model would undermine the power of the analysis. All features with reported significance for PD status that also appear significant in any model for a confounding variable are reported (Table S7).

Spearman’s correlation was conducted for clinical and dietary metadata variables using cor.test (stats R package). For gene and pathway feature levels, variables which showed an association (q ≤ 0.25) with PD donors in at least one multivariate analysis from MaAsLin2 are included. All correlations for a specific feature type between dietary or clinical variables categories are corrected for multiple comparisons together. To determine broad, high-level alterations in metabolic and functional microbial processes we performed pathway enrichment analysis on KEGG Orthology hierarchies utilizing a hypergeometric *P*-value test (Eq. 4) for PD vs. HC and PD vs. PC separately. Gene families with an association to disease status are defined by (q ≤ 0.25) in prior MaAsLin2 modeling and partitioned into enrichment and depletion:

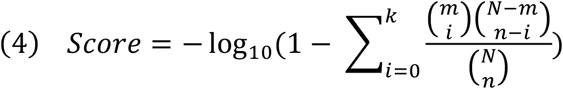

where *N* is the number of total features quantified, *n* is the total number of associated features, *m* is the number of detected features in the pathway, and *k* is the number of associated features in the pathway.

To identify the most robust metagenomic microbial feature associations with PD, we accessed metagenomics reads from two additional studies on Parkinson’s disease^20,52^ (NCBI BioProject accessions PRJEB17784 and PRJNA433459). Generalized linear mixed models were employed with MaAsLin2, testing only for PD status as a fixed effect with study of origin as a random effect. As a scale-invariant approach for quantifying feature effect sizes we implemented AUROC values with the pROC R package^53,54^. Values for features of each cohort were calculated using relative abundance profiles and confidence intervals were generated using permutations.

## RESULTS

### Subject Demographics Associate with Microbiome Composition

Stool samples from Parkinson’s Disease (PD), Population Controls (PC), and Household Controls (HC) were processed and analyzed for metagenomic (shotgun) sequencing (Fig. 1A). Biospecimens were collected from studies at the BioCollective (TBC) and Rush University Medical Center (RUMC), with the inclusion of HCs without PD allowing for analysis of a shared environment which is known to impact microbiome composition. Descriptive statistics for demographic metadata and PD clinical features are shown in Table S1. To assess metadata-explained variance in microbial community composition, we combined cohorts and conducted a PERMANOVA on various feature levels using the Aitchison distance, a linear measure of sample dissimilarity for compositional data^55^ (Fig. 1B; Table S2). We found several significant anthropomorphic, environmental, drug, and donor-group related associations. Notable environmental factors included household (R^2^ = 29.1 to 30.2 %) and cohort effects (R^2^ = 2.4 to 8.3 %), which independently explained a large percentage of variance. Disease status, donor group, and disease severity imparted significant effects on gut microbiome composition. General and PD-specific medication usage also appeared to explain gut microbiome variance and were tested for confounding potential (Table S7).

**FIG. 1.**
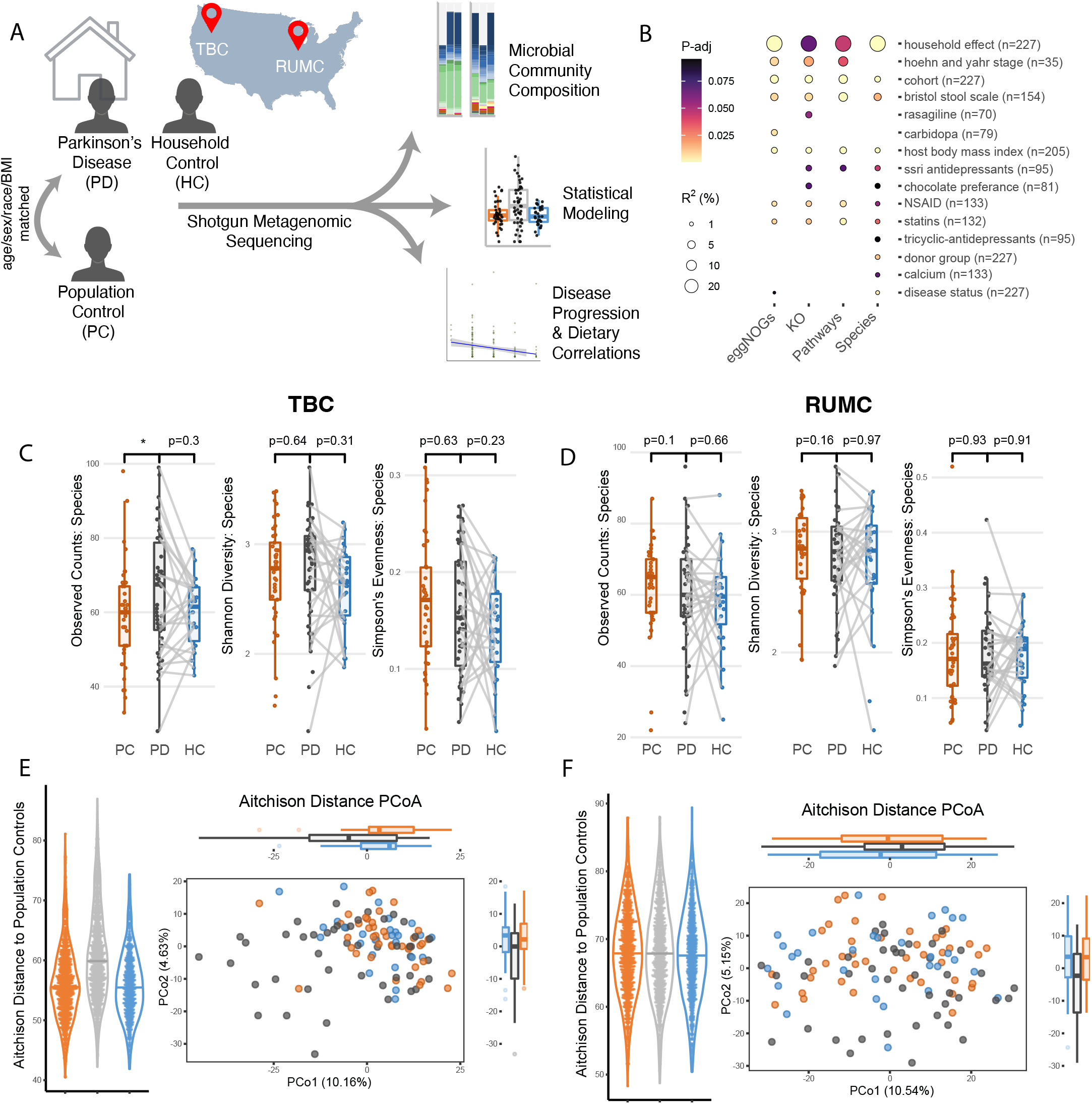
Parkinson’s disease microbial communities differ from healthy population and house-hold controls. **(A)**Schematic visualizing the donor groups and two distinct cohorts within our study. **(B)** Metadata explained variance across various feature levels using a PERMANOVA on the Aitchison distance. **(C-D)** Alpha diversity measures of observed species, Shannon diversity, and Simpson’s evenness on TBC **(C)** and RUMC **(D)** cohorts. **(E-F)** Relative abundance of species displayed by principal coordinates analysis (PCoA) of Aitchison distance on TBC **(E)** and RUMC **(F)** cohorts. Violin plots (left) show distance of samples within each group from samples from the population control group.

### Microbial Community Composition and Differentially Abundant Taxa

Diversity metrics provide a high-level assessment of microbial community composition. Alpha-diversity analyses revealed a subtle increase in unique taxa in PD subjects relative to PC in the TBC cohort (Fig. 1C), while no differences were observed in the RUMC samples (Fig. 1D). These findings are in agreement with previous studies showing a modest or no additional unique taxa in PD^33^. To better understand community structure and group variability, we separately calculated the Aitchison distance on species abundance within each cohort. PD status significantly explained a low percentage of variance in both TBC (R^2^ = 3.59 %) (Fig. 1E) and RUMC (R^2^ = 2.95 %) (Fig. 1F) cohorts, consistent with previous 16S rRNA gene sequencing studies^33^.

Differential abundance of individual taxa in disease states can inform potential microbiome influences on host physiology^56^. In joint modeling of cohorts, we observed several significant (q ≤ 0.1) and associated (q ≤ 0.25) taxonomic features in PD samples. Across both PD-PC and PD-HC comparisons, we observed enriched taxa in PD including associations with the Actinobacteria phylum (Fig. S2), and *Eisenbergiella tayi* and *Bifidobacterium bifidum* species (Fig. 2). Consistent PD-associated depletions included *Faecalibacterium* and *Roseburia* genera (Fig. S2), and *Faecalibacterium prausnitzii* species, known producers of short-chain fatty acids (Fig. 2). In accordance with previous studies, we observed PD-specific enrichment of *Akkermansia muciniphila* and *Ruthenibacterium lactatiformans* and depletions of *Eubacterium* species, though these differences were unique to either PD-HC or PD-PC comparisons (Fig. 2)^20,23,25,33^. These data show robust gut dysbiosis at the level of microbial abundance, but not taxonomic representation which is more uniform across study groups.

**FIG. 2.**
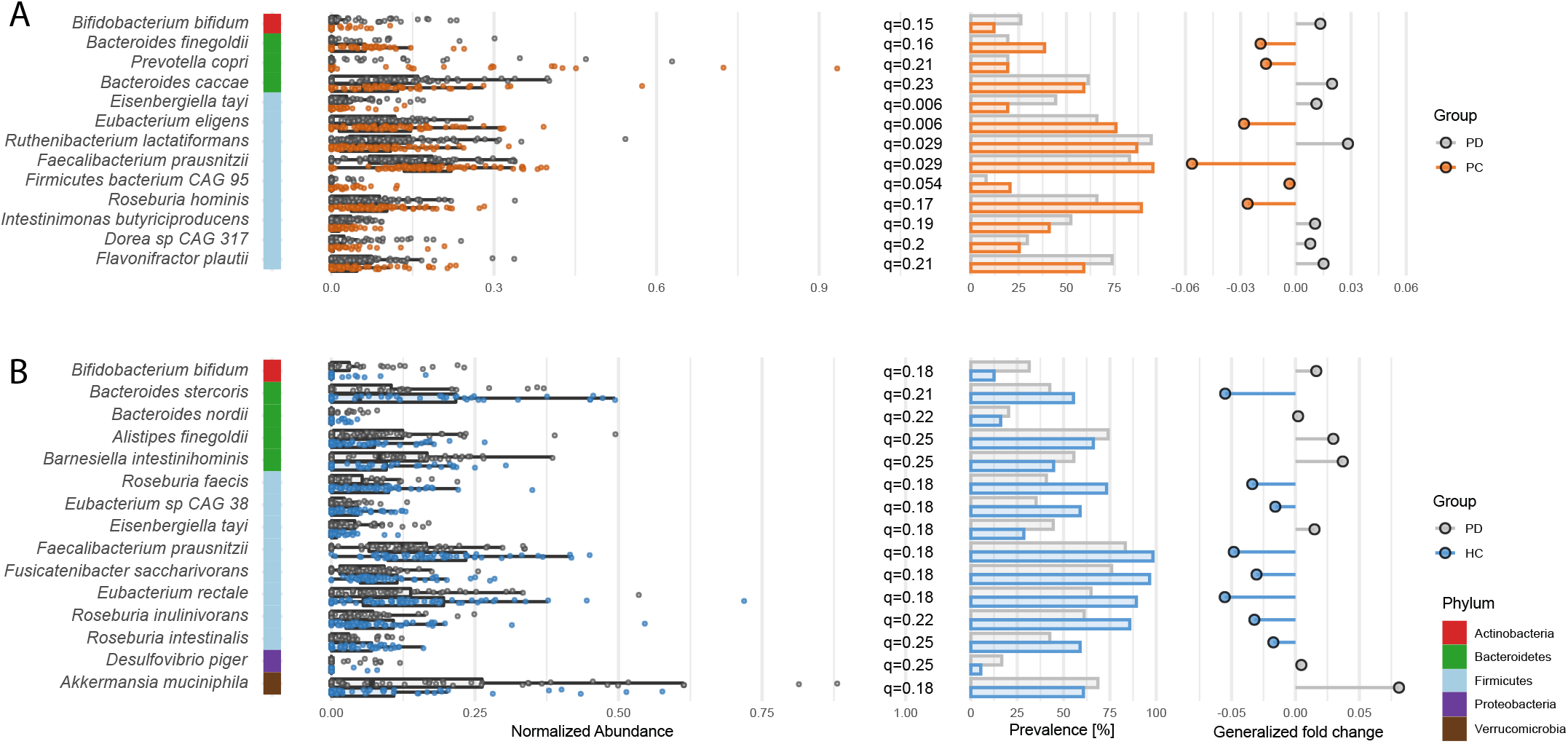
Parkinson’s disease patients have decreased abundance of putative beneficial anti-inflammatory short chain fatty acid-producing bacterial species. **(A, B)** Differentially abundant species determined through generalized linear models, displayed as relative abundance, prevalence (fraction each group with detection of taxa), and generalized or pseudo fold-change (average difference between groups across a range of quantiles). Comparison between **(A)** healthy population control (PC) samples and PD patients and **(B)** household controls (HC) and PD patients.

### Altered Metabolic Potential of the PD Metagenome

Microbial metabolites influence function of the metabolic, immune, and nervous systems in animals and humans^57–59^. We inferred the microbial metabolic potential of the PD microbiome through MetaCyc pathway abundance. Across comparisons between PD and both control groups, we observed significant increases in genes associated with pyruvate fermentation to acetone in the PD microbiome, and predicted depletions of CDP-diacylglycerol (CDP-DAG) biosynthesis steps I & II (Fig 3; Fig S3; Table S4). CDP-DAG is an essential intermediate in the production of phospholipids^60,61^. Downstream products of CDP-DAG can be integrated into bacterial membranes, and regulate cytoplasmic transport^62^. Pathway associations shared in both control group comparisons include 55 enrichments and 12 depletions in PD. The enrichments include synthesis and breakdown of fatty acids as well as biosynthesis of menaquinol, the reduced form of vitamin K2. Shared depletions in PD included synthesis of vitamins B1 and B2 (Fig 3A; Fig S3; Table S4).

**FIG. 3.**
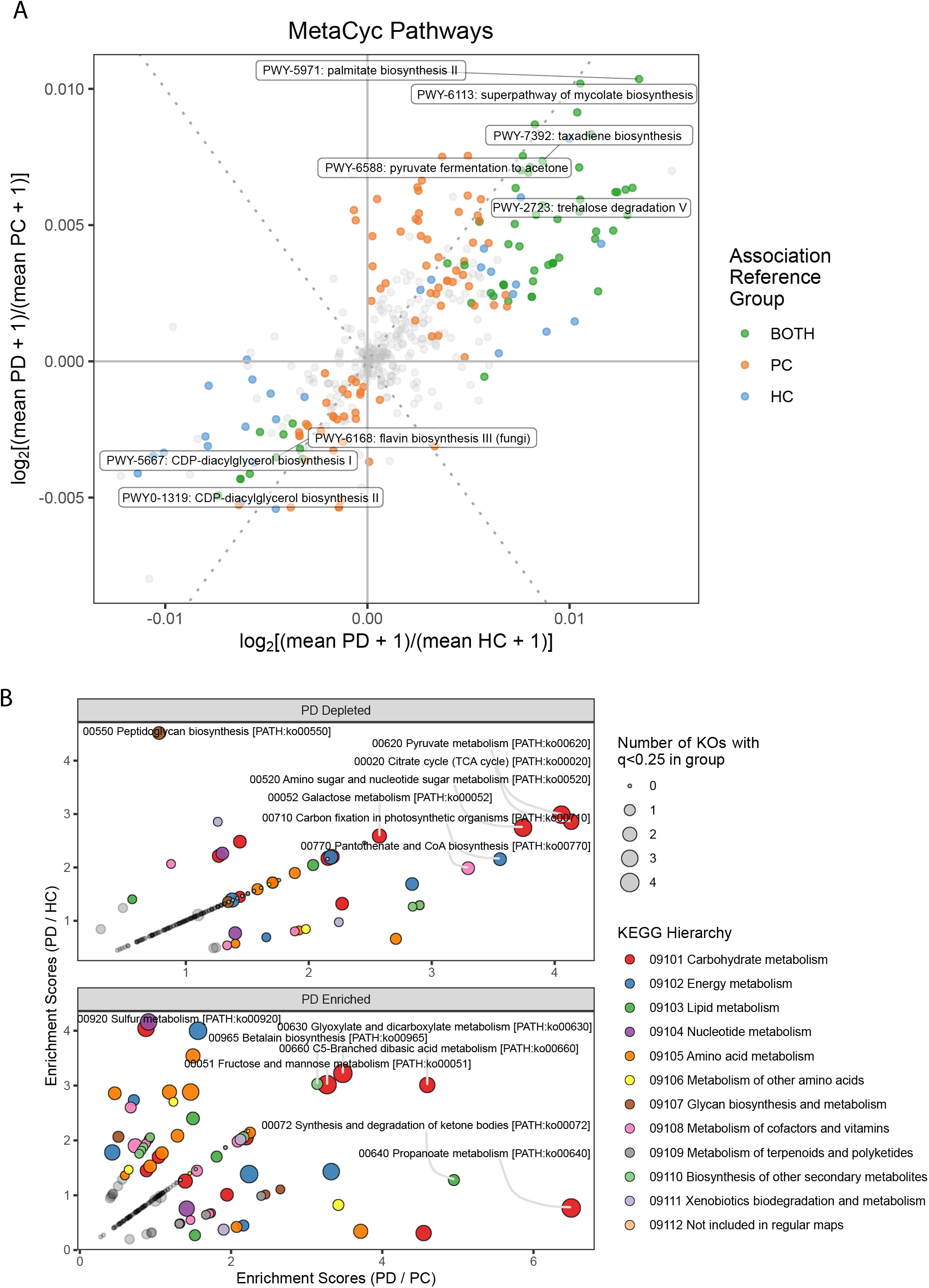
Microbial carbohydrate, amino acid, and lipid metabolism pathways are altered in Parkinson’s disease. )**(A)**Altered MetaCyc pathways in PD visualized as a scatterplot of log2 [(mean PD + 1)/ (mean HC + 1)] values by log2 [(mean PD + 1)/ (mean PC + 1)]. Color indicates feature associations determined using the general linear models previously described. Labels are provided for the top 8 most significant features based on the average q-value between analyses. **(B)** KEGG metabolism pathway enrichment analysis displayed by bubble plots.

For an unbiased synopsis of potential microbiome functions, we conducted pathway enrichment analysis of KEGG hierarchies which revealed notable shifts in carbohydrate, amino acid, and lipid metabolism (Fig. 3B). Genes for metabolic processes depleted in PD suggest alterations in carbohydrate metabolism with gene families related to glycolysis/gluconeogenesis, TCA cycle, and galactose metabolism. Enrichments across both PD-control comparisons included pathways for carbohydrate, amino acid, and lipid metabolism. These data are the first to highlight altered fatty acid metabolism in the gut microbiome of PD.

### Microbial Gene Signatures of PD

In addition to differences in functional metabolic pathways, we tested for alterations in the annotated genetic repertoire of the gut microbiome. Enzymes, KEGG Orthology (KO), Gene Ontology (GO), Pfams, and eggNOG gene families were constructed and assessed for disease associations.

Significantly enriched genes encoding enzymes shared across both PD comparisons included EC: 6.2.1.5 (succinate:CoA ligase (ADP-forming)) and EC: 1.3.5.2 ((S)-dihydroorotate:quinone oxidoreductase), while depletions included EC: 3.1.26.11 (tRNA 3’ endonuclease) and EC: 1.1.1.40 (malate dehydrogenase) (Table S3). KO analyses exclusively displayed shared significance for depletion including the lantibiotic transport system ATP-binding protein, and large subunit ribosomal protein L4. GO analyses revealed four shared enrichments and twelve depletions, including many genes involved in bacterial flagellar motility (Fig. S4). These alterations at the GO level are consistent with depletions in Pfams, largely of flagellar components, which are known to potently activate the immune system. For a complete summary of significant results for enzymes, KOs, GOs, Pfams, and eggNOGs, see Table S3.

Pathway enrichment analysis of KEGG hierarchies exhibited robust alterations in ABC transporters, with both enrichments and depletions in PD (Fig. S4C). Other disease state-enriched annotations include nucleotide and amino acid metabolism. Depleted hierarchies include cell growth, ribosomes, and multiple aspects of microbial motility and signaling (flagellar assembly, bacterial chemotaxis, and motility proteins), largely encoded within *Eubacterium* and *Roseburia*. This rich dataset may inform how biosynthetic functions harbored within the gut microbiome may influence disease status, a hypothesis that requires further investigation.

### Metagenomic Correlations with Disease Severity and Diet

Feature-wise associations with disease severity and dietary metadata were interrogated using Spearman’s rank-based correlations. We note 75 clinical and 252 dietary correlations across all metagenomic feature levels that are significantly altered. Of the features with the strongest and most significant correlations to clinical metadata (|ρ| ≤ 0.75 & q ≤ 0.1), we observed 9 distinguishing associations (Fig. 4A). MDS-UPDRS Part III scores positively associated with three KO gene families including anthranilate synthase component I (Fig. 4B). Curiously, loss of olfactory function, as captured by olfactory diagnosis and the UPSIT smell score, showed strong associations with microbial adenosine de-novo biosynthesis. Both correlations suggest that an increase of microbially produced adenosine is associated with an improved sense of smell.

**FIG. 4.**
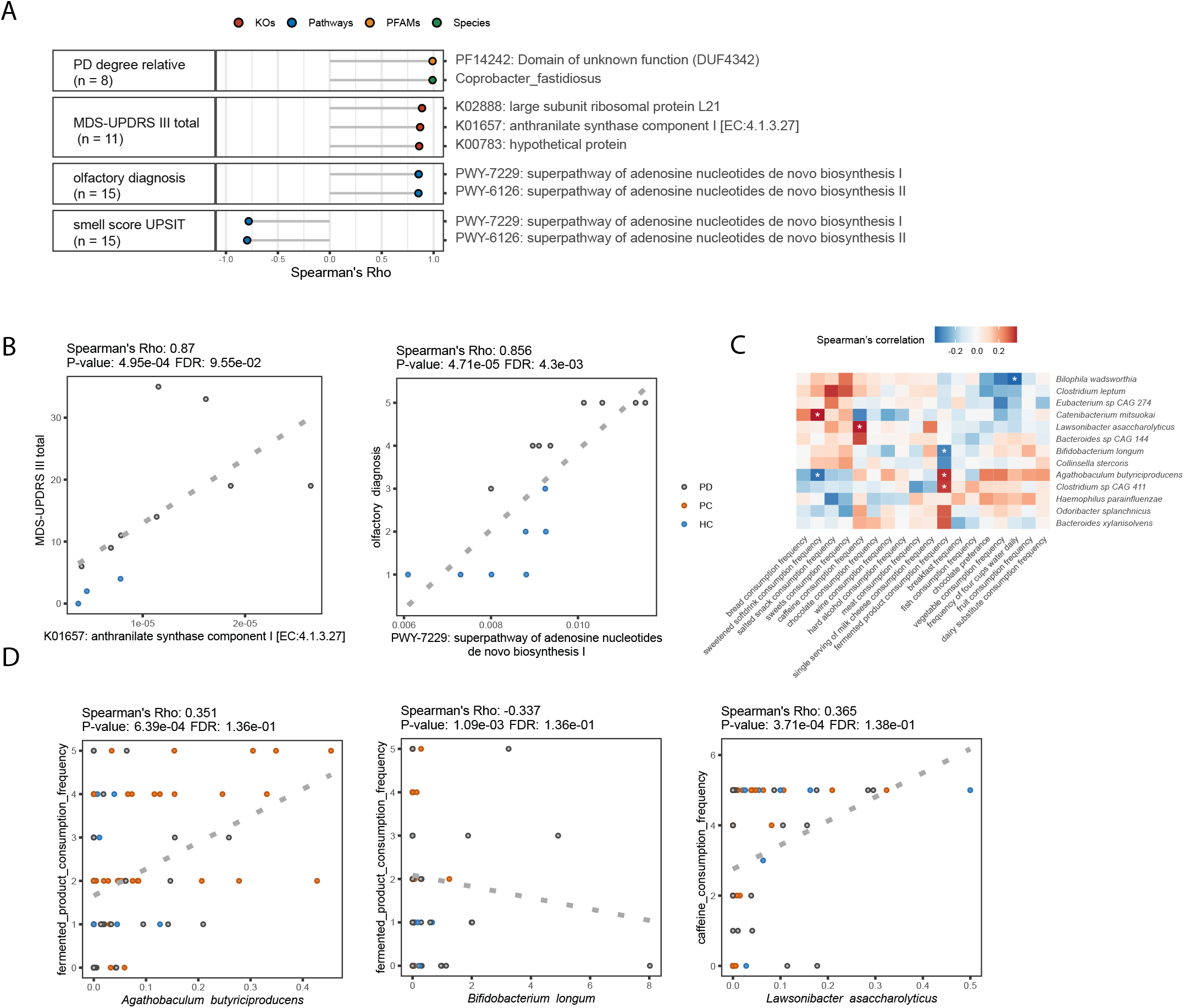
Dietary habits and disease severity correlate with metagenomic features. **(A)**Spearman’s correlations of metagenomic features and PD clinical metadata selecting for strongest associations (|ρ|≤ 0.75, FDR ≤ 0.1). Data points are colored by the feature level. **(B, D)** Scatterplots with linear regression for select correlations to PD clinical variables **(B)** and dietary habits **(D)**. **(C)** Heatmap of correlations between species and dietary survey variables. The top 13 species with the largest number of associations (q ≤ 0.25) were selected for visualization.

Several interesting correlations between taxa and dietary intake frequency were also captured. Fermented product consumption positively associated with *Agathobaculum butyriciproducens*, and negatively associated with *Bifidobacterium longum*, a microbe consistently enriched in PD patient microbiomes^20,52^(Fig. 4C,D). Additionally, we found that *Lawsonibacter asaccharolyticus* positively associated with caffeine consumption frequency, which has been proposed to be protective in PD^63–65^. The potential impact of this interesting correlation remains unknown. A comprehensive list of associations can be found in Table S5.

### Generalizable Microbial Signatures Across Four Independent PD Cohorts

Metagenomic features with generalizability for disease status across geographic locations and cohorts could provide tremendous value as potential biomarkers. Two previously assessed metagenomic cohorts of PD and controls, one from Bonn, Germany and another from Shanghai, China, were re-analyzed in parallel with the TBC and RUMC cohorts. Statistical analysis using all four cohorts revealed 12 species, 10 genera, 21 MetaCyc pathways, 232 Enzymes, 255 KOs, 541 GOs, 939 Pfams, and 1500 eggNOGs that significantly associated with disease status.

At the taxonomic level, we observed signatures concordant with previous 16S rRNA amplicon sequence analysis findings^19–30^. *R. lactatiformans, Gordonibacter pamelaeae, A. muciniphila*, and *Intestinimonas butyriciproducens* display a robust increase in PD. Consistently depleted taxa include *F. prausnitzii* and various *Roseburia* species (Fig. 5A; Fig S5A). Assessment of pathways related to production of cell membrane components showed an enrichment for lipopolysaccharide (LPS) and depletion in peptidoglycan biosynthesis, associated with Gram-negative and Gram-positive microbes, respectively (Fig. 5B). This may reflect a shift toward increased Gram-negative bacteria in PD, which is supported the observed increase in *Akkermansia muciniphila* and proportional decreases in *Faecalibacterium* and *Eubacterium* and *Roseburia* species.

**FIG. 5.**
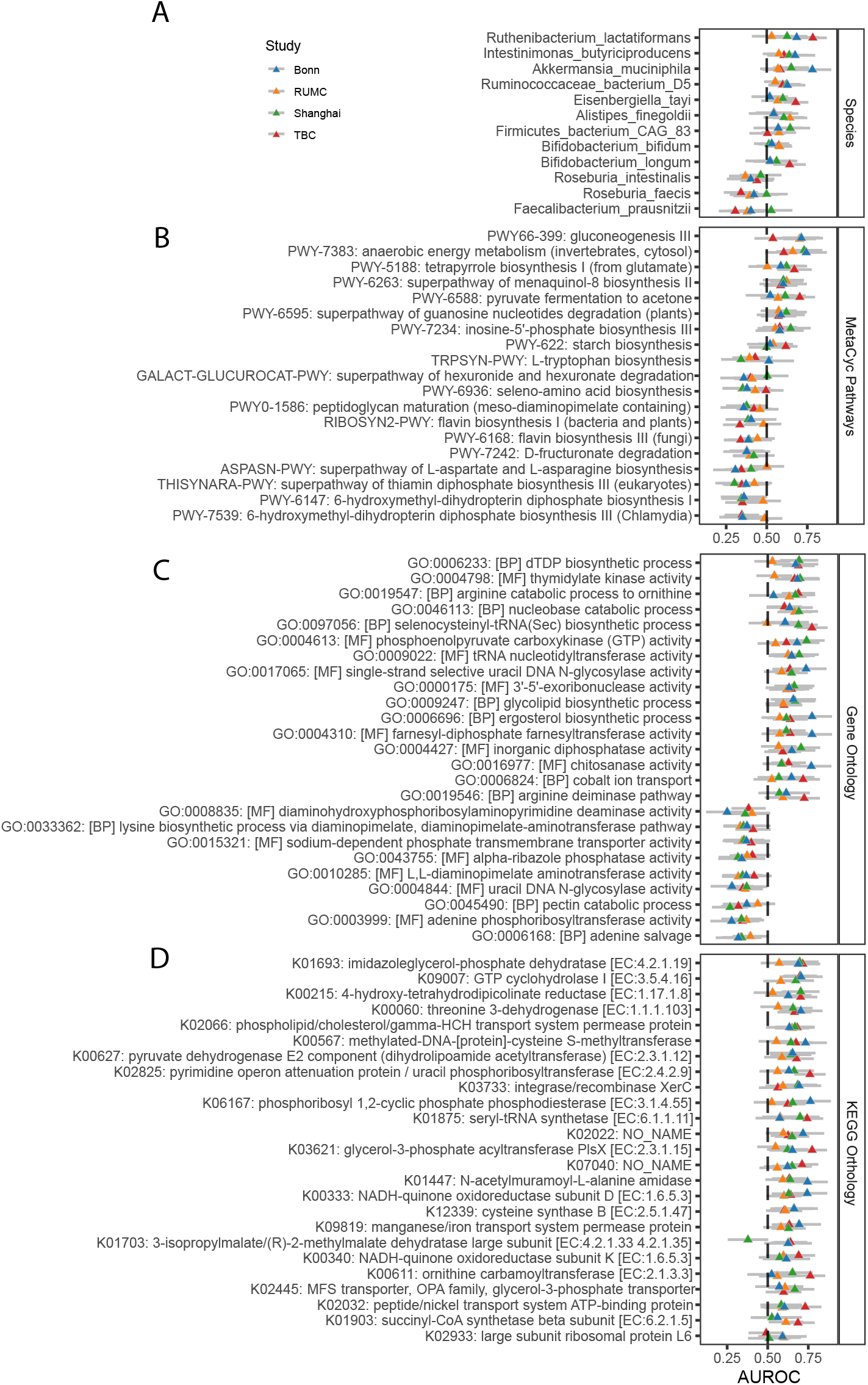
Meta-analysis of microbial features reveals generalizable Parkinson’s disease signatures across cohorts. **(A-D)** AUROC values and 95% confidence intervals per cohort for up to 25 significant features with the largest average absolute value of (AUROC-0.5) across all cohorts at the species **(A)**, MetaCyc pathway **(B)**, Gene Ontology **(C)**, and KEGG Orthology **(D)** levels.

Additional significant metabolic pathways and gene associations indicate signatures of oxidative stress and dysbiosis. Genes associated with response to oxidative stress included PD-specific enrichment of NAD-quinone oxidoreductase, SelR domains, indigoidine synthase A-like protein, and methionine sulfoxide reductase [EC 1.8.4.12] (Fig. 5C,D; Fig. S5).

Disrupted community signaling is inferred from signatures including numerous alterations in metal and ion transporters and permeases related to microbial defense mechanisms and quorum signaling (K09819: manganese/iron transport system permease protein, K02032: peptide/nickel transport system ATP-binding protein, etc.), proteins associated with biofilm formation and/or responses to hypoxic stress in biofilms (PF02567: phenazine biosynthesis-like protein, PF13277: YmdB-like protein), and PF07931: chloramphenicol phosphotransferase-like protein (Fig S5C).

Metabolism and biosynthesis of flavin and pterin compounds appear to be altered in PD. One of the strongest generalizable signatures was a PD enrichment in K09007: GTP cyclohydrolase I [EC:3.5.4.16] (Fig. 5D). Further, the MetaCyc pathway RIBOSYN2-PWY: flavin biosynthesis I (bacteria and plants) was considerably depleted in PD, along with PWY-6147: 6-hydroxymethyl-dihydropterin diphosphate biosynthesis I (Fig. 5B).

Collectively, despite differences in geographies (and associated diets), sequencing platform, and disease stage of study participants, analysis of this intercontinental sample set reveals that the PD microbiome displays robust and consistent signatures that differ from healthy controls.

Functional consequences or diagnostic application of these findings to motor performance, GI symptoms, or other aspects of PD remains to be determined.

## DISCUSSION

Microbiome changes at the taxonomic (e.g., genus, species) level have been described for various neuropsychiatric and neurodegenerative disorders, including PD. Herein, we employed shotgun metagenomics to reveal that the PD microbiome shows consistent shifts not only in microbial composition, but also alterations in functional metabolic processes related to cellular growth, and gene-level signatures, potentially indicative of a gut environment under stress. In our meta-analysis, we identify multiple taxonomic, metabolic, and gene-level microbial features with consistent changes across multiple PD cohorts. Recurring signatures include putative responses to oxidative stress and disruption of microbial community signaling. Enrichment of methionine sulfoxide reductase [EC 1.8.4.12] and SelR are strongly suggestive of an oxidative environment, since these enzymes reduce the oxidized form of methionine, which plays a significant role in reactive oxygen species (ROS) damage^66^. Additionally, indigoidine synthase A-like protein is a potent antioxidant and pterin and flavin are important co-factors in enzymatic redox reactions. It is tempting to speculate that enrichment of these microbial elements is a response to oxidative stress in the gut, potentially in response to the intestinal inflammation that has been reported in PD^25,67^. Interestingly, oxidative stress has previously been associated with alpha-synuclein misfolding, a contributor to PD pathophysiology^68,69^.

Additional gene-level signatures suggestive of a dysbiotic microbial environment include a depletion in the quorum signaling GO pathway (GO:0009372), robust depletion of flagellar genes, and increases in transporters with involvement in microbial defense mechanisms.

Curiously, depletion of flagellar genes in the microbiome has been associated with fibrosis severity in patients with non-alcoholic fatty liver disease^70^, and type 2 diabetes^71^. Additionally, loss of flagella has been tied to an increase in cellular growth rate^72^, possibly related to immune evasion. Flagella are highly immunogenic features targeted by both innate and adaptive host immune systems^73^. A possible explanation for the observed depletion of flagella may be increased targeting of flagellated bacteria by innate or adaptive immunity, a testable hypothesis with mucosal immune profiling of gut biopsies from PD patients.

Our data provide novel insight into gut microbial metabolic processes in PD patients, but several questions remain. A primary limitation of our study is the sparsity of metadata collection across cohorts by questionnaires about medication usage, limiting our ability to account for the impact of drug treatment in our modeling. As with all cross-sectional sampling, no cause-and-effect associations can be inferred from this study. Longitudinal studies incorporating individuals prior to onset of PD symptoms (i.e., prodromal cohorts), capturing newly diagnosed or treatment-naïve patients would be invaluable for determining directionality of gut microbiome perturbations and PD status or severity. While gut bacteria have been shown to modulate motor performance, intestinal transit, neuroinflammation, a-synuclein pathology and neurodegeneration in PD mouse models^74,75^, the contribution of the microbiome to outcomes in human PD remains speculative.

Together, our data suggest the PD microbiome is defined by modest alterations in community composition, but robust dysregulation in carbohydrate, lipid, and amino acid metabolism, an enrichment of gene-level signatures for responses to oxidative stress, and depletion of gene-level signatures of cellular growth, microbial motility, and disrupted community signaling. This study therefore presents novel generalizable disease associations that may lead to a better understanding of how the gut microbiome may influence, or be influenced by, PD symptoms and lifestyles.

## Supporting information

Supplementary Table 1: Clinical variables

Supplementary Table 1: Demographic variables

Supplementary Table 1: PD medication usage

## ACKNOWLEDGMENT

The authors thank Yvette Garcia-Flores for administrative and technical assistance, and Gabriella Sanzo and Alex Yerkan for assistance with patient recruitment and data collection. We thank Dr. Viviana Gradinaru, Dr. Catherine Oikonomou, and members of the Mazmanian laboratory for insightful evaluation of the manuscript. This research was funded in part by the Department of Defense (PD160030) to A.K., and S.K.M., Aligning Science Across Parkinson’s (ASAP-020495) through the Michael J. Fox Foundation for Parkinson’s Research (MJFF) to S.K.M., and the MJFF (Grant #15780) to S.K.M. For the purpose of open access, the author has applied a CC BY 4.0 public copyright license to all Author Accepted Manuscripts arising from this submission.

## AUTHOR’S ROLES

Design, J.C.B., M.C., A.K., S.K.M.; execution, J.C.B., L.A.V.M., D.A.H., P.A.E., Z.Z., D.L., G.H., G.A.; analysis, J.C.B., G.S., J.O., D.J.H., J.W.B., R.K., S.K.M.; writing, J.C.B., S.K.M.; editing, all authors.

## FINANCIAL DISCLOSURES OF ALL AUTHORS (for the preceding 12 months)

S.K.M. declares financial interests unrelated to this work in Axial Therapeutics, Nuanced Health, and Seed Health.

## DATA SHARING

Metagenomic sequencing data are available via Qiita study IDs 14476 and 12975 and EBI-ENA accessions ERP138197 and ERP138199 for TBC and RUMC cohorts, respectively. Detailed scripts to reproduce all analysis and data visualization are available at: https://github.com/jboktor/PD_Metagenomic_Analysis.git. In addition, we have made an interactive shiny R app available via the link above for exploration of metagenomic features of interest in our datasets (Fig. S6).

## TABLE AND FIGURE LEGENDS

**FIG.1 Supplement.**
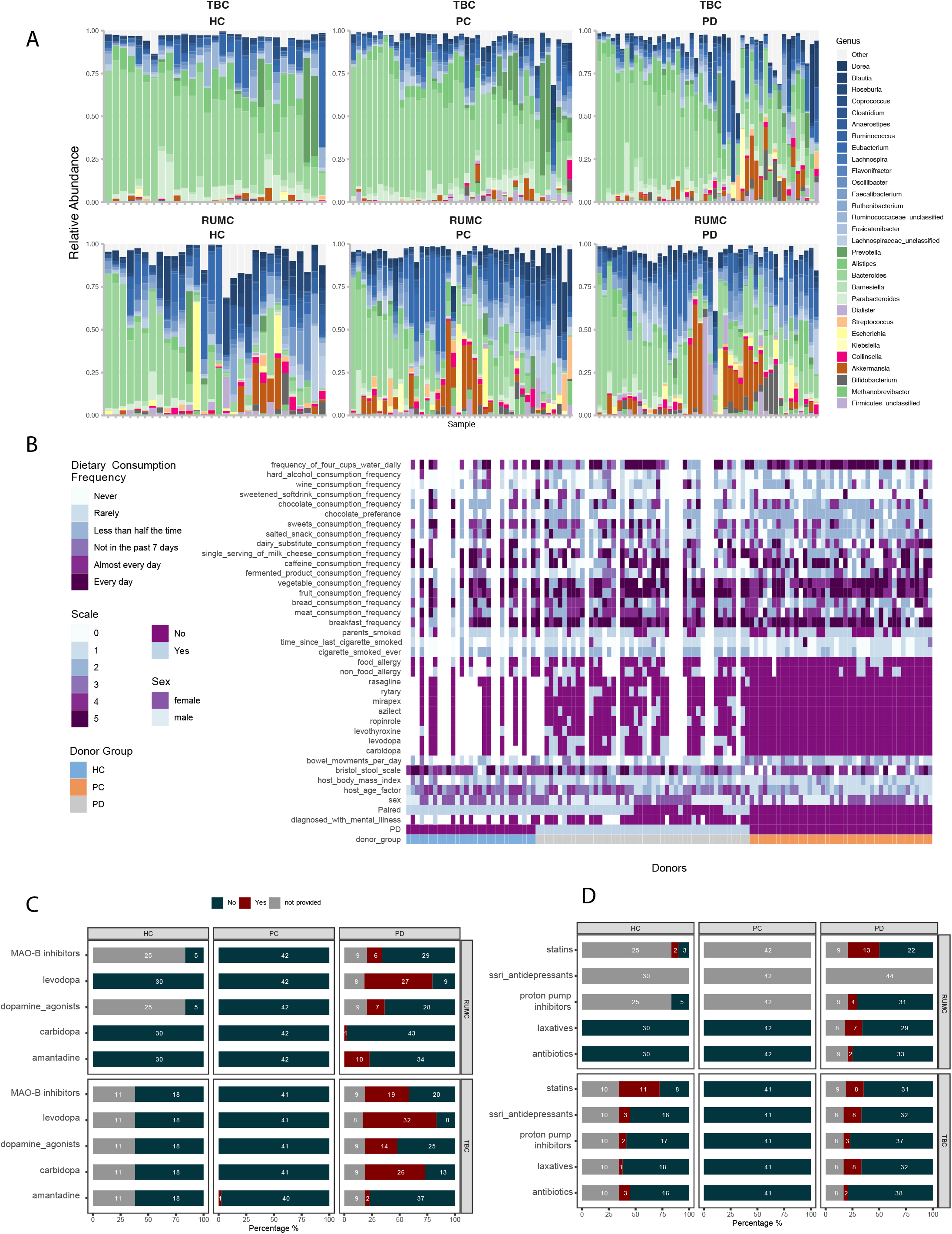
**(A)** Top 30 most abundant bacterial genera colored by gradient within the order level for both TBC (top) and RUMC (bottom) samples. **(B)** TBC survey responses displayed as a heatmap. **(C-D)** Stacked bar plots displaying responses for PD medications **(C)** and other known potentially confounding factors **(D)** from both cohorts.

**FIG.2 Supplement.**
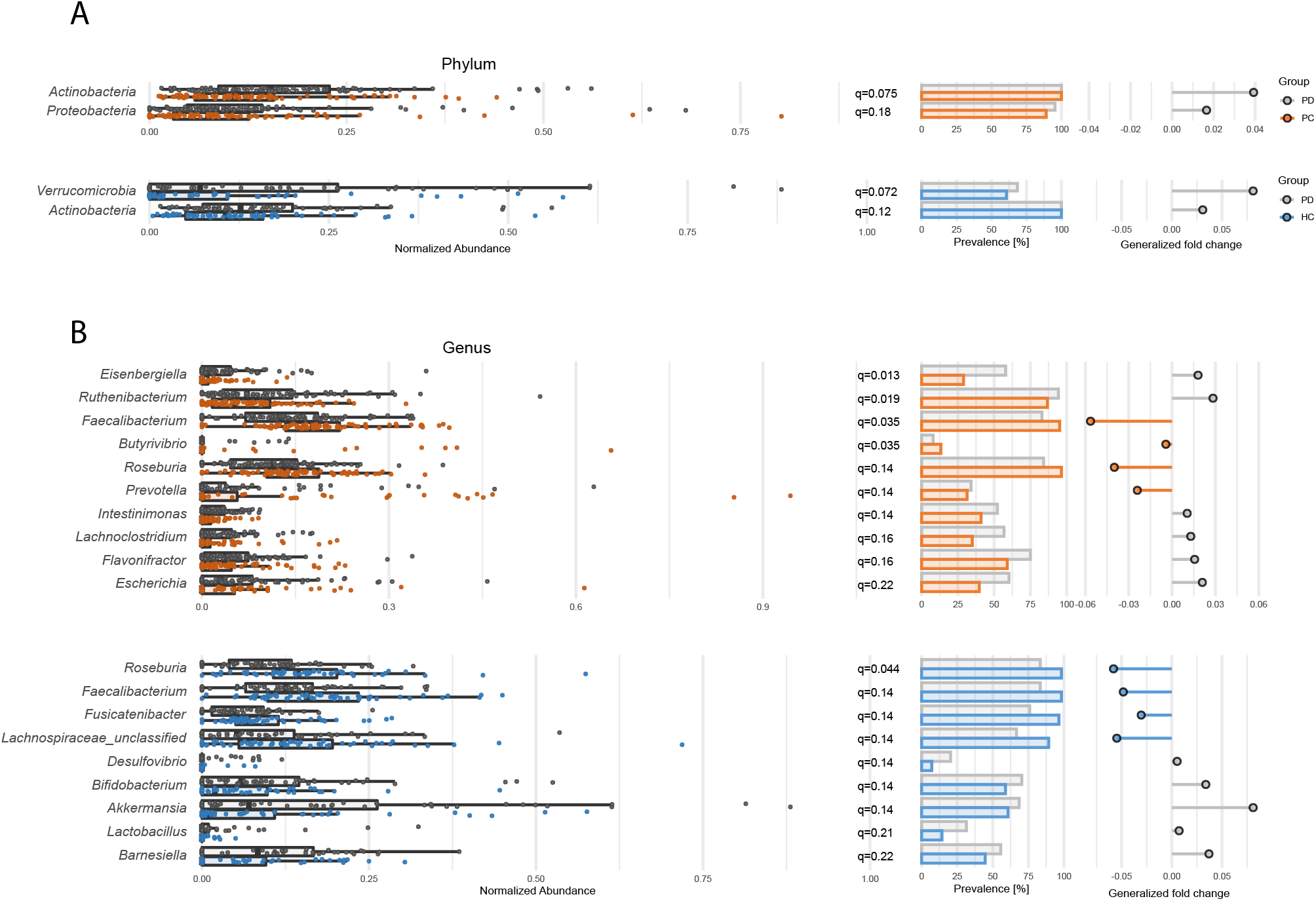
**(A, B)** Differentially abundant bacterial phyla and genera determined through general linear models are displayed by relative abundance, prevalence, and generalized or pseudo fold-change. Models are as previously described.

**FIG.3 Supplement.**
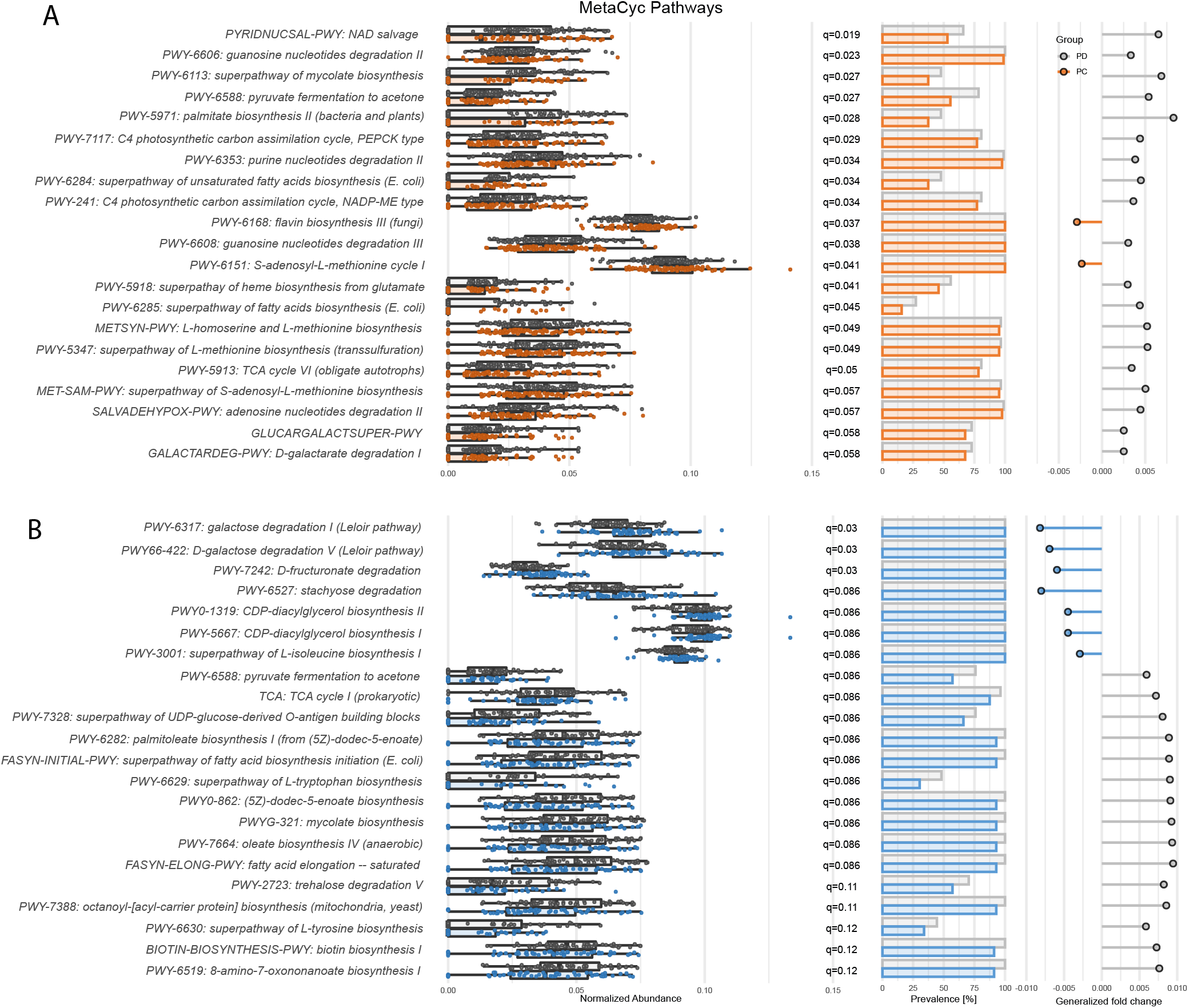
Differentially abundant MetaCyc pathways between **(A)** healthy population control (PC) samples and PD participants and **(B)** household controls (HC) and PD participants.

**FIG.4 Supplement.**
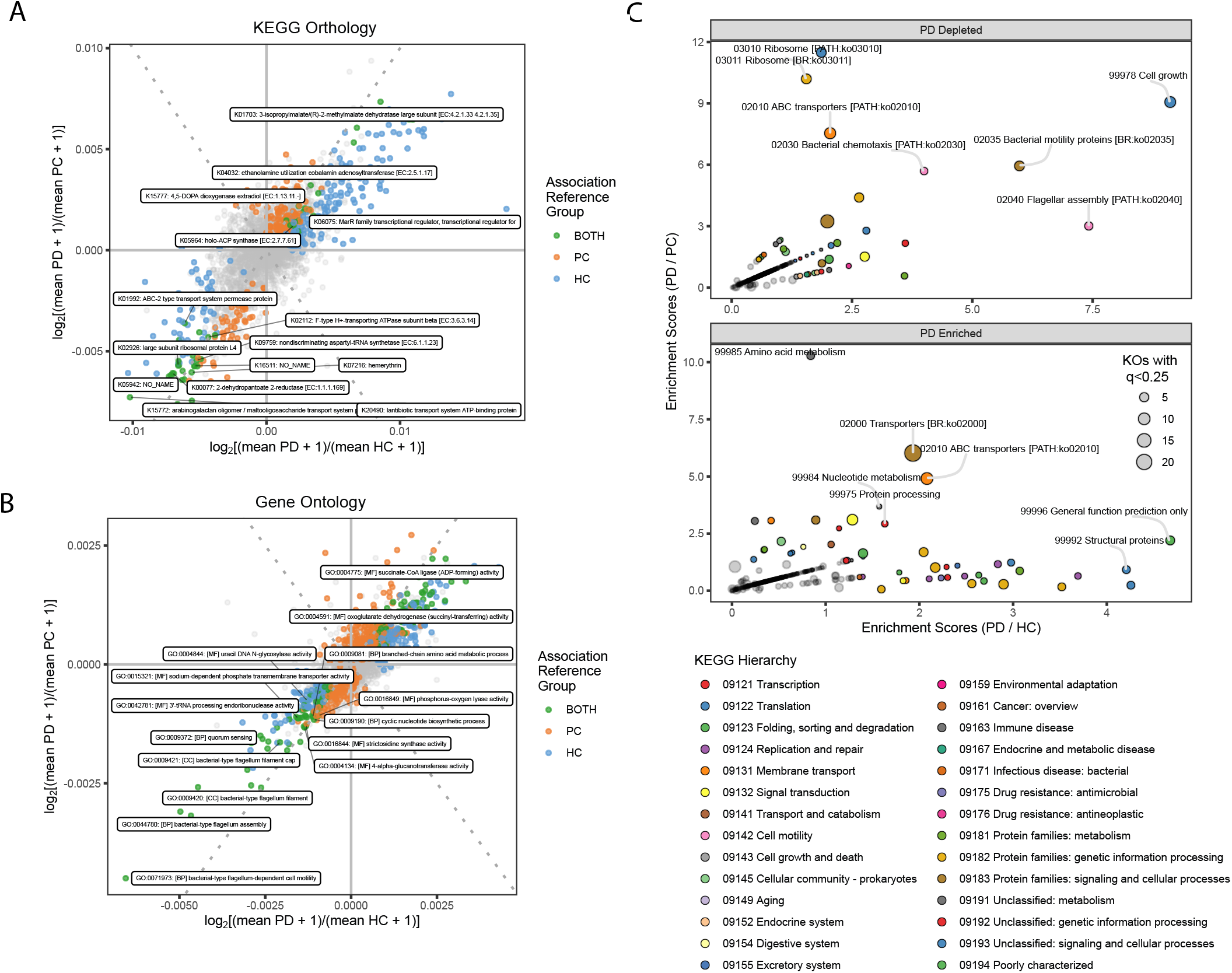
Gene-level analysis reveals signatures of intestinal stress. **(A-B)** Scatterplots visualizing features of interest for KEGG Orthology **(A)** and Gene-Ontology **(B)** features. **(C)** Bubble plots of enrichment for all other categories of KEGG annotation.

**FIG.5 Supplement.**
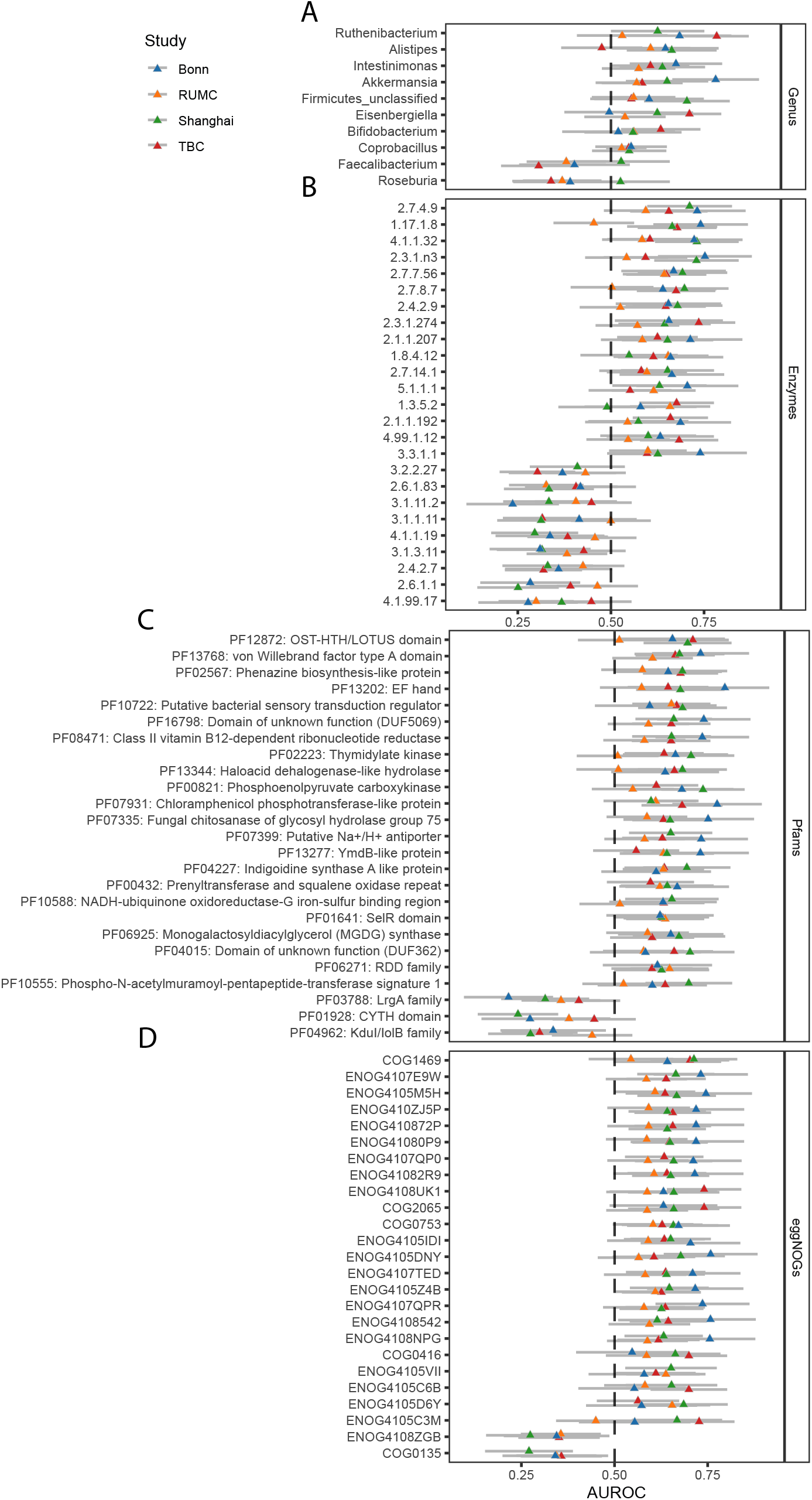
**(A-D)** AUROC values and 95% confidence intervals per cohort for up to 25 features with the largest average absolute value of (AUROC-0.5) across all cohorts at the genus **(A)**, enzyme **(B)**, Pfam **(C)** and eggNOG **(D)** levels.

**FIG.6 Supplement.**
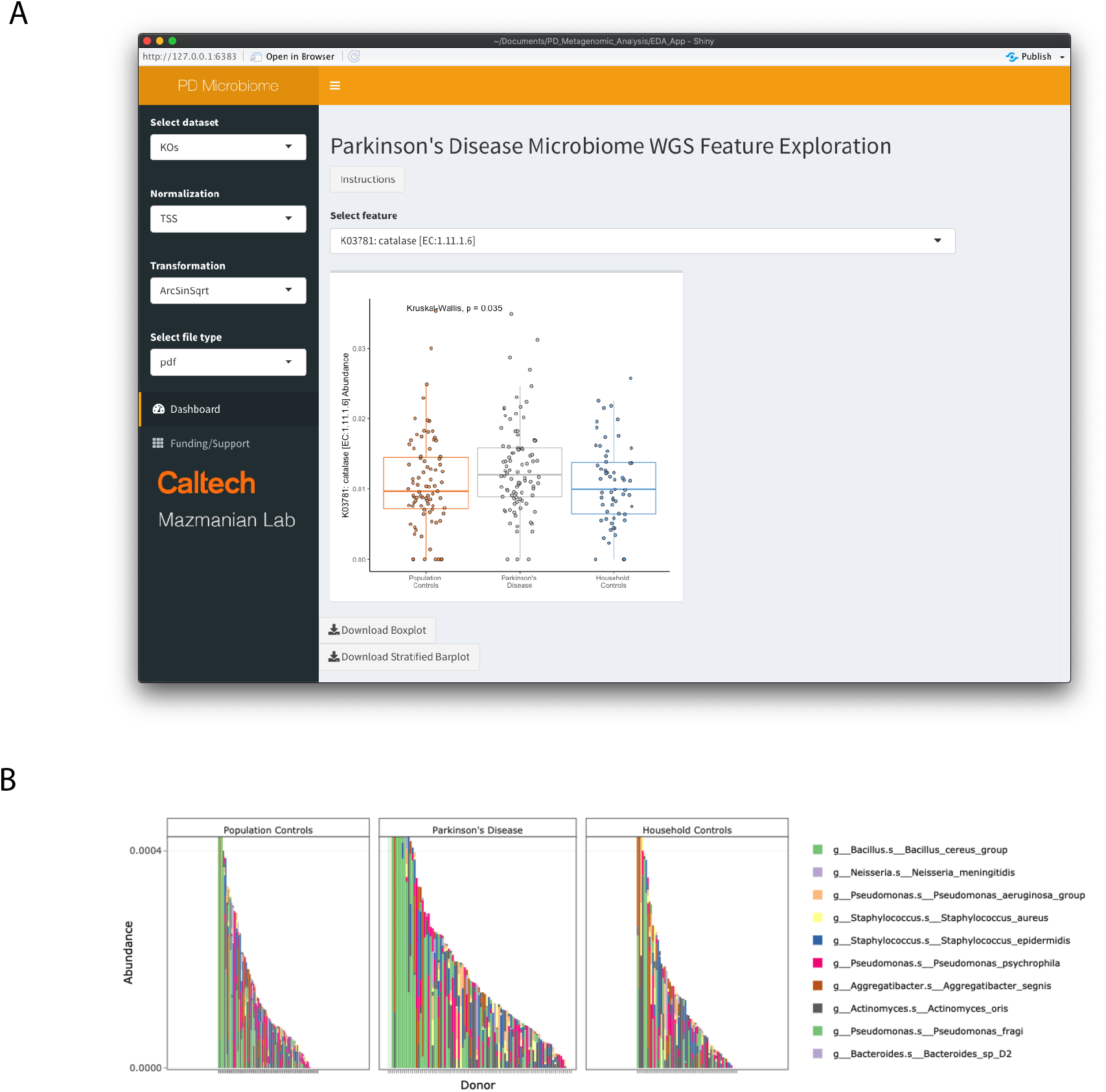
**(A)** Shiny application interface for exploration of metagenomic features in PD. **(B)** Example of interactive downloadable stratified abundance table.

## SUPPLEMENTARY TABLES

Table S1: Sample Demographics Table S2: PERMANOVA statistics

Table S3: Differentially Abundant Features

Table S4: Pathway Enrichment Analysis

Table S5: Spearman’s Correlation with Dietary Metadata

Table S6: Spearman’s Correlation with Clinical Metadata

Table S7: Confounder Estimation

Table S8: Feature AUROC and Meta-Analysis

## SUPPLEMENTARY METHODS

### Automated Self-Administered 24-Hour Recall

The PD and HC participants completed the Automated Self-Administered 24-hour (ASA24®) recall. The ASA24® recall is a validated and reliable web-based tool developed by the National Cancer Institute to capture 24-hour dietary intake^44,76,77^. The PD and HC participants reported each food item that they consumed in the last 24 hours using the gold standard, automated multiple-pass method (AMPM)^77^. The Computer-Assisted Self-Interviewing (CASI) methodology is used to guide the respondent through multiple steps of recalls that include reporting each meal or snack or any other time that food or beverage was consumed, a comprehensive list of foods and drinks consumed, and finally a detail step that includes quantity of food consumed, any forgotten foods, and a final review. The criterion validity of the ASA24® recall is supported by high agreement (∼80%) with traditional interviewer-administered recalls and comparable energy intake estimates between ASA24® recalls and AMPM in healthy men and women^76,78^. The ASA24® recall was used to identify dietary intake consistency to determine its influence on gut microbiota and secondary outcome measurements.

### Vioscreen Dietary Questionnaire

The PD and HC participants completed the Vioscreen Food Frequency Questionnaire, an adult-validated, self-administered, web-based dietary assessment tool^45^. This questionnaire captures 19 food components.

### University of Pennsylvania Smell Identification Test

The PD and HC participants completed the University of Pennsylvania Smell Identification Test (UPSIT)^46^. The UPSIT is a self-administered clinical test of olfactory function (i.e, evaluating changes in smell) that uses microencapsulated odorants which are released by scratching standardized odor-impregnated test booklets.

### Sleep Behavior Disorder and Sleep Disturbance Questionnaires

The PD and HC participants completed the idiopathic rapid eye movement (REM) sleep behavior disorder (RBD) single-question screen (RBD1Q). The RBD1Q is a screening tool for diagnosis of REM Sleep Behavior Disorder, which is an important risk factor for PD^47^. The RBD1Q questionnaire consists of a single question, answered “yes” or “no,” as follows: “Have you ever been told, or suspected yourself, that you seem to ‘act out your dreams’ while asleep (for example, punching, flailing your arms in the air, making running movements, etc.)?” Additionally, the PD and HC participants completed the Pittsburgh Sleep Quality Index (PSQI)^48^, which is a self-rated questionnaire which assesses sleep quality and disturbances over a 1-month time interval.

### Munich ChronoType Questionnaire

The PD and HC participants completed the Munich ChronoType Questionnaire (MCTQ) which contains 60 questions pertaining to sleep and activity times such as: bedtime, length of time to fall asleep, time of awakening, and use of alarm clock on workdays and work-free days^49^. All questions are asked separately for work and for work-free days. Chronotype is estimated as the midpoint of sleep on work-free days minus half of the difference between sleep duration on work-free days and average sleep duration of the week to control for sleep debt (Midpoint of Sleep on Work-Free days, Sleep-Corrected, MSFsc)^79^. The classification of the MCTQ can be extreme early, normal or extreme late^80^. The MCTQ was also used to calculate social jet lag and sleep debt^79^. Social jet lag (SJL) is the difference between the timing of the midpoint of sleep on work days and work-free days^81^, and is considered significant if greater than two hours^82^. Sleep debt is the difference between average sleep duration for the week and sleep duration during work days^81^.

### Patient-Reported Outcomes Measurements Information System Gastrointestinal Symptom Scale

The PD and HC participants completed the validated National Institutes of Health (NIH) Patient-Reported Outcomes Measurements Information System (PROMIS) gastrointestinal symptom scale. The NIH PROMIS uses eight GI symptom scales that are used for clinical care and research across the full range of GI disorders^50^. Only four GI symptoms scales were measured across time for this study. Belly pain (six questions), bowel incontinence (four questions), constipation (9 questions), and gas & bloating (12 questions) were reported by both the PD and HC participants. Higher scores denoted more GI symptoms. Lower scores denoted less GI symptoms. Scores range from 20 (low) to 80 (high). A score of 50 denotes the general population average.

## REFERENCES

1. Marras, C. et al. Prevalence of Parkinson’s disease across North America. npj Parkinson’s Disease 4, 1–7 (2018).

2. Postuma, R. B. et al. MDS clinical diagnostic criteria for Parkinson’s disease. Movement Disorders 30, 1591–1601 (2015).

3. Mhyre, T. R., Boyd, J. T., Hamill, R. W. & Maguire-Zeiss, K. A. Parkinson’s Disease. Subcell Biochem 65, 389–455 (2012).

4. Lesage, S. & Brice, A. Parkinson’s disease: from monogenic forms to genetic susceptibility factors. Human Molecular Genetics 18, R48–R59 (2009).

5. Nonnekes, J., Post, B., Tetrud, J. W., Langston, J. W. & Bloem, B. R. MPTP-induced parkinsonism: an historical case series. The Lancet Neurology 17, 300–301 (2018).

6. Mantri, S., Duda, J. E. & Morley, J. F. Early and Accurate Identification of Parkinson Disease Among US Veterans. Fed Pract 36, S18–S23 (2019).

7. Goldman, S. et al. Polychlorinated Biphenyls (PCBs) and Parkinson’s Disease (PD): Effect Modification by Membrane Transporter Variants (S32.004). Neurology 86, (2016).

8. Moustafa, A. A. et al. Motor symptoms in Parkinson’s disease: A unified framework. Neuroscience & Biobehavioral Reviews 68, 727–740 (2016).

9. Durcan, R. et al. Prevalence and duration of non-motor symptoms in prodromal Parkinson’s disease. Eur J Neurol 26, 979–985 (2019).

10. Postuma, R. B. et al. Identifying prodromal Parkinson’s disease: Pre-Motor disorders in Parkinson’s disease. Movement Disorders 27, 617–626 (2012).

11. Schapira, A. H. V., Chaudhuri, K. R. & Jenner, P. Non-motor features of Parkinson disease. Nat Rev Neurosci 18, 435–450 (2017).

12. Park, A. & Stacy, M. Non-motor symptoms in Parkinson’s disease. J Neurol 256, 293–298 (2009).

13. Round, J. L. & Mazmanian, S. K. The gut microbiome shapes intestinal immune responses during health and disease. Nat Rev Immunol 9, 313–323 (2009).

14. Mazmanian, S. K., Liu, C. H., Tzianabos, A. O. & Kasper, D. L. An Immunomodulatory Molecule of Symbiotic Bacteria Directs Maturation of the Host Immune System. Cell 122, 107–118 (2005).

15. Visconti, A. et al. Interplay between the human gut microbiome and host metabolism. Nat Commun 10, 4505 (2019).

16. Fung, T. C., Olson, C. A. & Hsiao, E. Y. Interactions between the microbiota, immune and nervous systems in health and disease. Nat Neurosci 20, 145–155 (2017).

17. Vuong, H. E. et al. The maternal microbiome modulates fetal neurodevelopment in mice. Nature 586, 281–286 (2020).

18. Lozupone, C. A., Stombaugh, J. I., Gordon, J. I., Jansson, J. K. & Knight, R. Diversity, stability and resilience of the human gut microbiota. Nature 489, 220–230 (2012).

19. Aho, V. T. E. et al. Gut microbiota in Parkinson’s disease: Temporal stability and relations to disease progression. EBioMedicine 44, 691–707 (2019).

20. Bedarf, J. R. et al. Functional implications of microbial and viral gut metagenome changes in early stage L-DOPA-naïve Parkinson’s disease patients. Genome Med 9, 39 (2017).

21. Hasegawa, S. et al. Intestinal Dysbiosis and Lowered Serum Lipopolysaccharide-Binding Protein in Parkinson’s Disease. PLOS ONE 10, e0142164 (2015).

22. Heintz-Buschart, A. et al. The nasal and gut microbiome in Parkinson’s disease and idiopathic rapid eye movement sleep behavior disorder. Movement Disorders 33, 88–98 (2018).

23. Hill-Burns, E. M. et al. Parkinson’s disease and Parkinson’s disease medications have distinct signatures of the gut microbiome. Movement Disorders 32, 739–749 (2017).

24. Hopfner, F. et al. Gut microbiota in Parkinson disease in a northern German cohort. Brain Research 1667, 41–45 (2017).

25. Keshavarzian, A. et al. Colonic bacterial composition in Parkinson’s disease. Movement Disorders 30, 1351–1360 (2015).

26. Li, W. et al. Structural changes of gut microbiota in Parkinson’s disease and its correlation with clinical features. Sci. China Life Sci. 60, 1223–1233 (2017).

27. Li, F. et al. Alteration of the fecal microbiota in North-Eastern Han Chinese population with sporadic Parkinson’s disease. Neuroscience Letters 707, 134297 (2019).

28. Lin, A. et al. Gut microbiota in patients with Parkinson’s disease in southern China. Parkinsonism & Related Disorders 53, 82–88 (2018).

29. Petrov, V. A. et al. Analysis of Gut Microbiota in Patients with Parkinson’s Disease. Bull Exp Biol Med 162, 734–737 (2017).

30. Unger, M. M. et al. Short chain fatty acids and gut microbiota differ between patients with Parkinson’s disease and age-matched controls. Parkinsonism & Related Disorders 32, 66–72 (2016).

31. Beghini, F. et al. Integrating taxonomic, functional, and strain-level profiling of diverse microbial communities with bioBakery 3. eLife 10, e65088 (2021).

32. Sharpton, T. J. An introduction to the analysis of shotgun metagenomic data. Front Plant Sci 5, 209 (2014).

33. Romano, S. et al. Meta-analysis of the Parkinson’s disease gut microbiome suggests alterations linked to intestinal inflammation. npj Parkinsons Dis. 7, 1–13 (2021).

34. Wallen, Z. D. et al. Characterizing dysbiosis of gut microbiome in PD: evidence for overabundance of opportunistic pathogens. npj Parkinsons Dis. 6, 1–12 (2020).

35. Cirstea, M. S. et al. Microbiota Composition and Metabolism Are Associated With Gut Function in Parkinson’s Disease. Movement Disorders 35, 1208–1217 (2020).

36. Baldini, F. et al. Parkinson’s disease-associated alterations of the gut microbiome predict disease-relevant changes in metabolic functions. BMC Biol 18, 62 (2020).

37. Hertel, J. et al. Integrated Analyses of Microbiome and Longitudinal Metabolome Data Reveal Microbial-Host Interactions on Sulfur Metabolism in Parkinson’s Disease. Cell Rep 29, 1767-1777.e8 (2019).

38. Rosario, D. et al. Systematic analysis of gut microbiome reveals the role of bacterial folate and homocysteine metabolism in Parkinson’s disease. Cell Reports 34, 108807 (2021).

39. Movement Disorder Society Task Force on Rating Scales for Parkinson’s Disease. The Unified Parkinson’s Disease Rating Scale (UPDRS): status and recommendations. Mov Disord 18, 738–750 (2003).

40. Goetz, C. G. et al. Movement Disorder Society Task Force report on the Hoehn and Yahr staging scale: status and recommendations. Mov Disord 19, 1020–1028 (2004).

41. Hughes, A. J., Daniel, S. E., Kilford, L. & Lees, A. J. Accuracy of clinical diagnosis of idiopathic Parkinson’s disease: a clinico-pathological study of 100 cases. J Neurol Neurosurg Psychiatry 55, 181–184 (1992).

42. Couch, R. D. et al. The approach to sample acquisition and its impact on the derived human fecal microbiome and VOC metabolome. PLoS One 8, e81163 (2013).

43. Zreloff, Z. J., Lange, D., Vernon, S. D., Carlin, M. R. & Cano, R. de J. Accelerating Gut Microbiome Research with Robust Sample Collection. (2021) doi:10.20944/preprints202101.0047.v1.

44. Subar, A. F. et al. The Automated Self-Administered 24-hour dietary recall (ASA24): a resource for researchers, clinicians, and educators from the National Cancer Institute. J Acad Nutr Diet 112, 1134–1137 (2012).

45. Kristal, A. R. et al. Evaluation of web-based, self-administered, graphical food frequency questionnaire. J Acad Nutr Diet 114, 613–621 (2014).

46. Doty, R. L., Shaman, P., Kimmelman, C. P. & Dann, M. S. University of Pennsylvania Smell Identification Test: a rapid quantitative olfactory function test for the clinic. Laryngoscope 94, 176–178 (1984).

47. Postuma, R. B. et al. A single-question screen for rapid eye movement sleep behavior disorder: a multicenter validation study. Mov Disord 27, 913–916 (2012).

48. Buysse, D. J., Reynolds, C. F., Monk, T. H., Berman, S. R. & Kupfer, D. J. The Pittsburgh Sleep Quality Index: a new instrument for psychiatric practice and research. Psychiatry Res 28, 193–213 (1989).

49. Roenneberg, T., Wirz-Justice, A. & Merrow, M. Life between clocks: daily temporal patterns of human chronotypes. J Biol Rhythms 18, 80–90 (2003).

50. Spiegel, B. M. R. et al. Development of the NIH Patient-Reported Outcomes Measurement Information System (PROMIS) gastrointestinal symptom scales. Am J Gastroenterol 109, 1804–1814 (2014).

51. Mallick, H. et al. Multivariable Association Discovery in Population-scale Meta-omics Studies. 2021.01.20.427420 https://www.biorxiv.org/content/10.1101/2021.01.20.427420v1 (2021) doi:10.1101/2021.01.20.427420.

52. Qian, Y. et al. Gut metagenomics-derived genes as potential biomarkers of Parkinson’s disease. Brain 143, 2474–2489 (2020).

53. Robin, X. et al. pROC: an open-source package for R and S+ to analyze and compare ROC curves. BMC Bioinformatics 12, 77 (2011).

54. Wirbel, J. et al. Meta-analysis of fecal metagenomes reveals global microbial signatures that are specific for colorectal cancer. Nat Med 25, 679–689 (2019).

55. Gloor, G. B., Macklaim, J. M., Pawlowsky-Glahn, V. & Egozcue, J. J. Microbiome Datasets Are Compositional: And This Is Not Optional. Front Microbiol 8, 2224 (2017).

56. Matheoud, D. et al. Intestinal infection triggers Parkinson’s disease-like symptoms in Pink1-/-mice. Nature 571, 565–569 (2019).

57. Kim, C. H. Immune regulation by microbiome metabolites. Immunology 154, 220–229 (2018).

58. Zhao, S. et al. Dietary fructose feeds hepatic lipogenesis via microbiota-derived acetate. Nature 579, 586–591 (2020).

59. Needham, B. D. et al. A gut-derived metabolite alters brain activity and anxiety behaviour in mice. Nature 602, 647–653 (2022).

60. Dowhan, W. CDP-diacylglycerol synthase of microorganisms. Biochimica et Biophysica Acta (BBA) - Lipids and Lipid Metabolism 1348, 157–165 (1997).

61. Sohlenkamp, C. & Geiger, O. Bacterial membrane lipids: diversity in structures and pathways. FEMS Microbiology Reviews 40, 133–159 (2016).

62. Slavetinsky, C., Kuhn, S. & Peschel, A. Bacterial aminoacyl phospholipids - Biosynthesis and role in basic cellular processes and pathogenicity. Biochim Biophys Acta Mol Cell Biol Lipids 1862, 1310–1318 (2017).

63. Nefzger, M. D., Quadfasel, F. A. & Karl, V. C. A Retrospective Study of Smoking in Parkinson’s Disease. American Journal of Epidemiology 88, 149–158 (1968).

64. Ascherio, A. et al. Caffeine, postmenopausal estrogen, and risk of Parkinson’s disease. Neurology 60, 790–795 (2003).

65. Lee, K.-W. et al. Neuroprotective and Anti-inflammatory Properties of a Coffee Component in the MPTP Model of Parkinson’s Disease. Neurotherapeutics 10, 143–153 (2013).

66. Singh, V. K. & Moskovitz, J. 2003. Multiple methionine sulfoxide reductase genes in Staphylococcus aureus: expression of activity and roles in tolerance of oxidative stress. Microbiology 149, 2739–2747.

67. Devos, D. et al. Colonic inflammation in Parkinson’s disease. Neurobiology of Disease 50, 42–48 (2013).

68. Scudamore, O. & Ciossek, T. Increased Oxidative Stress Exacerbates α-Synuclein Aggregation In Vivo. Journal of Neuropathology & Experimental Neurology 77, 443–453 (2018).

69. Park, J.-H. et al. Alpha-synuclein-induced mitochondrial dysfunction is mediated via a sirtuin 3-dependent pathway. Molecular Neurodegeneration 15, 5 (2020).

70. Schwimmer, J. B. et al. Microbiome Signatures Associated With Steatohepatitis and Moderate to Severe Fibrosis in Children With Nonalcoholic Fatty Liver Disease. Gastroenterology 157, 1109–1122 (2019).

71. Qin, J. et al. A metagenome-wide association study of gut microbiota in type 2 diabetes. Nature 490, 55–60 (2012).

72. Korem, T. et al. Growth dynamics of gut microbiota in health and disease inferred from single metagenomic samples. Science 349, 1101–1106 (2015).

73. Cullender, T. C. et al. Innate and Adaptive Immunity Interact to Quench Microbiome Flagellar Motility in the Gut. Cell Host & Microbe 14, 571–581 (2013).

74. Sampson, T. R. et al. Gut Microbiota Regulate Motor Deficits and Neuroinflammation in a Model of Parkinson’s Disease. Cell 167, 1469-1480.e12 (2016).

75. Sampson, T. R. et al. A gut bacterial amyloid promotes α-synuclein aggregation and motor impairment in mice. eLife 9, e53111 (2020).

76. Kirkpatrick, S. I. et al. Performance of the Automated Self-Administered 24-hour Recall relative to a measure of true intakes and to an interviewer-administered 24-h recall. Am J Clin Nutr 100, 233–240 (2014).

77. Moshfegh, A. J. et al. The US Department of Agriculture Automated Multiple-Pass Method reduces bias in the collection of energy intakes. Am J Clin Nutr 88, 324–332 (2008).

78. Thompson, F. E. et al. Comparison of Interviewer-Administered and Automated Self-Administered 24-Hour Dietary Recalls in 3 Diverse Integrated Health Systems. Am J Epidemiol 181, 970–978 (2015).

79. Kantermann, T., Sung, H. & Burgess, H. J. Comparing the Morningness-Eveningness Questionnaire and Munich ChronoType Questionnaire to the Dim Light Melatonin Onset. J Biol Rhythms 30, 449–453 (2015).

80. Fischer, D., Lombardi, D. A., Marucci-Wellman, H. & Roenneberg, T. Chronotypes in the US -Influence of age and sex. PLoS One 12, e0178782 (2017).

81. Wittmann, M., Dinich, J., Merrow, M. & Roenneberg, T. Social jetlag: misalignment of biological and social time. Chronobiol Int 23, 497–509 (2006).

82. Roenneberg, T., Allebrandt, K. V., Merrow, M. & Vetter, C. Social jetlag and obesity. Curr Biol 22, 939–943 (2012).

