## Supplementary Table 1: Clinical variables for "Integrated multi-cohort analysis of the Parkinson’s disease gut metagenome"

| Characteristic | N | N = 88[1] |
|---|---|---|
| disease_duration | 40 | 7.0 (5.0, 11.0) |
| age_of_onset | 40 | 58 (51, 61) |
| bristol_stool_scale | 72 | 3.00 (2.00, 4.00) |
| updrs_total | 35 | 17 (11, 22) |
| mds_updrs_3_total | 8 | 16 (10, 22) |
| mds_updrs_survey_total | 14 | 30 (17, 40) |
| hy_stage | 32 | 2.0000 (2.0000, 2.0000) |
| smell_score_upsit | 8 | 19.0 (14.5, 23.5) |
| olfactory_diagnosis | 8 | 4.50 (4.00, 5.00) |

[1] Median (IQR)
