## Supplementary Table 1: Demographic variables for "Integrated multi-cohort analysis of the Parkinson’s disease gut metagenome"

| Characteristic | N | Rush-HC, N = 30[1] | Rush-PC, N = 42[1] | Rush-PD, N = 42[1] | TBC-HC, N = 26[1] | TBC-PC, N = 41[1] | TBC-PD, N = 46[1] | p-value[2] |
|---|---|---|---|---|---|---|---|---|
| host_age | 225 | 62 (59, 70) | 66 (65, 69) | 62 (59, 68) | 70 (59, 73) | 56 (48, 62) | 68 (57, 73) | <0.001 |
| sex | 227 | | | | | | | 0.069 |
| female | | 18 (60%) | 15 (36%) | 12 (29%) | 14 (54%) | 21 (51%) | 21 (46%) | |
| male | | 12 (40%) | 27 (64%) | 30 (71%) | 12 (46%) | 20 (49%) | 25 (54%) | |
| host_body_mass_index | 205 | 27 (22, 32) | 34 (32, 40) | 27 (22, 29) | 25 (24, 28) | 24 (21, 25) | 24 (22, 26) | <0.001 |
| total_reads | 227 | 22,578,266 (19,139,010, 24,040,706) | 25,179,790 (21,169,508, 27,678,346) | 19,726,516 (16,047,950, 23,503,010) | 9,235,316 (8,818,786, 11,453,143) | 10,989,657 (8,714,403, 13,727,201) | 11,642,414 (9,111,935, 13,946,721) | <0.001 |
| bristol_stool_scale | 154 | 4.00 (3.00, 6.00) | NA (NA, NA) | 3.00 (2.00, 3.75) | 4.00 (4.00, 5.00) | 4.00 (4.00, 5.00) | 4.00 (2.25, 4.00) | 0.001 |

[1] Median (IQR); n (%)

[2] Kruskal-Wallis rank sum test; Pearson's Chi-squared test
