## Supplementary Table 1: PD medication usage for "Integrated multi-cohort analysis of the Parkinson’s disease gut metagenome"

| Characteristic | N | N = 88[1] |
| --- | --- | --- |
| levodopa | 72 | 55 (76%) |
| carbidopa | 79 | 26 (33%) |
| dopamine_agonists | 70 | 18 (26%) |
| MAO_B_inhibitors | 70 | 24 (34%) |
| rasagiline | 70 | 10 (14%) |
| selegiline | 70 | 15 (21%) |

[1] n (%)
